# Agrammatism and paragrammatism: a cortical double dissociation revealed by lesion-symptom mapping

**DOI:** 10.1101/2020.03.24.006759

**Authors:** William Matchin, Alexandra Basilakos, Brielle C. Stark, Dirk-Bart den Ouden, Julius Fridriksson, Gregory Hickok

## Abstract

The fundamental distinction of grammatical deficits in aphasia, agrammatism and paragrammatism, was made over a century ago. However, the extent to which the agrammatism/paragrammatism distinction exists independently of differences in speech fluency has not clearly been investigated. Despite much research on agrammatism, the lesion correlates of paragrammatism are essentially unknown. Lesion-symptom mapping was used to investigate the degree to which the lesion correlates of agrammatism and paragrammatism overlap or dissociate. Four expert raters assessed videos of 53 right-handed patients with aphasia following chronic left hemisphere stroke retelling the Cinderella story. Consensus discussion determined each subject’s classification with respect to grammatical deficits as Agrammatic, Paragrammatic, Both, or No Grammatical Deficit. Each subject’s lesion was manually drawn on a high-resolution MRI and warped to standard space for group analyses. Lesion-symptom mapping analyses were performed in NiiStat including lesion volume as a covariate. Secondary analyses included speech rate (words per minute) as an additional covariate. Region of interest analyses identified a double dissociation between these syndromes: damage to Broca’s area was significantly associated with agrammatism, p = 0.001 (but not paragrammatism, p = 0.930), while damage to the left posterior superior and middle temporal gyri was significantly associated with paragrammatism, p < 0.001 (but not agrammatism, p = 0.873). The same results obtained when regressing out the effect of speech rate, and non-overlapping lesion distributions between the syndromes were confirmed by uncorrected whole brain analyses. Our results support a fundamental distinction between agrammatism and paragrammatism.

## Introduction

Kleist (1914) noted two kinds of syntactic disturbances in the speech of patients with aphasia: agrammatism and paragrammatism. Agrammatism is described as the simplification of grammatical structure and omission of function words and morphemes. For example, Goodglass (1993) reports this example of agrammatic speech:

> Examiner: What brought you to the hospital?
>
> Patient: Yeah… Wednesday, … Paul and dad… Hospital… yeah… doctors, two…an’ teeth.

Paragrammatism, by contrast, is the misuse of grammar not attributable to an overall reduction of grammatical morphemes or simplification of syntactic structure. Kleist noted the prodigious output of paragrammatic patients, leading to “confused sentence monsters”. Butterworth and Howard (1987) provide examples of paragrammatic speech:

> “And I want everything to be so talk.”
>
> “She was handled to look at the books a bit.”
>
> “I’m very want it.”
>
> “Isn’t look very dear, is it?”
>
> “But it’s silly, aren’t they?”

Agrammatism has received extensive treatment in the literature and is widely known by both researchers and clinicians. By contrast, there is very little research on paragrammatism, and clinical practitioners are often unaware of its existence at all, leaving potential grammatical deficits in aphasia undiagnosed. Agrammatism is associated with non-fluent aphasia, primarily Broca’s aphasia (Saffran et al., 1989; Damasio, 1992; Goodglass, 1993; Thompson et al., 1997), while paragrammatism is associated with fluent aphasia, such as Wernicke’s and conduction aphasia (Goodglass *et al*., 1993). While recent lesion-behavior mapping studies have associated agrammatic speech with damage to primarily frontal structures, particularly Broca’s area (Sapolsky *et al*., 2010; Wilson *et al*., 2010; den Ouden *et al*., 2019), only a few case studies suggest an association of paragrammatism with posterior temporal-parietal damage (Yagata *et al*., 2017; Wilson *et al*., 2018*a*; *b*). Consistent with this, fluent aphasia is mostly associated with posterior temporal-parietal lesions (Buchsbaum *et al*., 2011; Ogar *et al*., 2011; Fridriksson *et al*., 2014; Yourganov *et al*., 2016). However, there have yet to be any large-scale lesion-symptom mapping studies of paragrammatic deficits *per se*. Therefore, in the present work, we sought to identify the lesion distribution associated with paragrammatism as distinct from agrammatism using voxel-based lesion-symptom mapping in a large cohort of people with chronic aphasia following left-hemisphere stroke.

An important issue concerning research into syntactic deficits in aphasia is the question of how syntax is organized in the healthy brain. While prominent models of syntax in the brain posit a primary syntactic function to different subregions of the inferior frontal gyrus (Friederici 2017; Hagoort 2014), Matchin and Hickok (2019) recently hypothesized a new model of syntax in the brain, with two primary cortical zones responsible for distinct aspects of syntactic processing. The first is a hierarchical lexical-syntactic system in the left posterior middle temporal gyrus (pMTG), including the ventral bank of the superior temporal sulcus (STS). This system underlies hierarchical structures necessary for conceptual-semantic processing that are critical for both comprehension and production of language (Everaert et al., 2015). The second is a linear morpho-syntactic system in the left inferior frontal gyrus, pars triangularis (IFGtri). This system underlies linear morpho-syntactic relations (such as word order and affixation rules) that are critical for producing speech through the vocal tract, which is primarily a serial system (Idsardi & Raimey, 2013; Berwick & Chomsky, 2016). However, the IFGtri is not strictly necessary for comprehension, as comprehension does not require the explicit computation of serial relations, which are provided by the incoming sequence of morphological inputs (Lewis & Vasishth, 2005).

Thus, Matchin & Hickok predict that damage to pMTG impairs both syntactic comprehension and production, whereas damage to IFGtri primarily impairs syntactic production (with potential deficits for comprehension of complex sentences due to confounding effects of impaired working memory resources; Rogalsky & Hickok, 2011; Matchin, 2017). However, the model also predicts asymmetric production deficits following from damage to these two systems: paragrammatic production deficits result from damage to the hierarchical lexical-syntactic system in the pMTG, whereas agrammatic production deficits result from damage to the linear morpho-syntactic system in the IFGtri. This is the prediction we tested in the current study.

We identified patients’ grammatical deficits through a perceptual rating of patients’ speech samples. According to the hypothesis presented in Matchin & Hickok (2019), we expected a double dissociation: paragrammatism would be associated with damage to the pMTG (and not IFGtri) while agrammatism would be associated with damage to IFGtri (and not the pMTG). Because neuroimaging studies have also identified syntactic effects in neighboring tissue (see Matchin & Hickok, 2019 for a review), and the difficulty of straightforwardly interpreting the localization in lesion-symptom mapping (Wilson, 2016), we created broader regions of interest (ROIs) encompassing these regions in order to maximize our ability to detect effects.

Importantly, the agrammatism/paragrammatism distinction has been argued to be governed by fluency rather than by a fundamental underlying grammatical distinction. From this perspective, both patient groups may have the same grammatical deficit, but agrammatic patients’ speech production problems prevent them from producing much output at all, whereas paragrammatic patients’ intact speech production allows them to produce voluminous disordered output (Heeschen, 1985; Heeschen and Kolk, 1988; Kolk and Heeschen, 1992). Therefore, in secondary analyses we included a speech rate measure (words per minute) as a covariate to account for potential confounding effects of overall speech fluency.

A note on notation: throughout this manuscript, we express the general concepts discussed in the literature of agrammatism (and agrammatic speech) and paragrammatism (and paragrammatic speech) using regular typeface. We express the corresponding perceptual classification of these concepts as applied to our subject groups using small capital typeface, i.e. agrammatism/agrammatic and paragrammatism/paragrammatic.

## Procedure

### Subjects

We initially analyzed connected speech samples from 100 people (28 women) with chronic post-stroke aphasia, which were collected as part of larger studies at the University of South Carolina and the Medical University of South Carolina. All subjects were recruited through local advertisement. They provided informed consent to participate in this study, which was approved by the Institutional Review Boards at the University of South Carolina and the Medical University of South Carolina. All subjects had a single ischemic stroke to the left hemisphere at least six months prior to study inclusion and were also pre-morbidly right handed (self-disclosed). On average, subjects were 59.2 years old at time of testing (±11.15 yrs), were 47.22 months post-stroke (± 47.78 months), had 14.89 years of education (± 2.41 years) and a Western Aphasia Battery - Revised [WAB-R] aphasia quotient [AQ] of 58.70 (± 21.30) (Kertesz, 2007). Out of the initial 100 subjects, the proportions of the following aphasia types were included: 19 with anomia, 50 with Broca’s aphasia, 17 with conduction aphasia, six with global aphasia, two with transcortical motor aphasia, and six with Wernicke’s aphasia.

### Classification of grammatical deficits

Clinical rating scales have been developed for agrammatism, such as the Boston Diagnostic Aphasia Examination (BDAE) grammatical form measure (Goodglass *et al*., 2000). However, no modern evaluative tools exist for classification of paragrammatism. We therefore relied on qualitative consensus assessments of expert raters, using the traditional Kleist (1914) criteria for agrammatism and paragrammatism. To elicit speech, we followed the story retelling protocol from AphasiaBank (MacWhinney *et al*., 2011). Subjects first reviewed a picture book of the Cinderella story (text omitted), then the book was removed and they were asked to retell the Cinderella story in their own words. Video recordings were made, lasting from a few seconds to more than five minutes. Subjects were instructed to use content from the book and their own recollection. Four expert raters (authors WM, AB, BS, DdO), blind to the subject’s aphasia type, psychological assessment scores, and lesions, individually watched these recordings and rated each subject as agrammatic, paragrammatic, no grammatical deficit, (ngd, i.e., no specific grammatical deficit, with possible presence of other aphasic deficits) or cannot rate based on informal perceptual judgments. Raters were allowed to listen to each video as many times as they liked. Following individual rating of each of the 100 subjects, a discussion was held with all four experts to resolve disagreements about each case and to determine a consensus rating. Patients exhibiting features of both agrammatism and paragrammatism were classified as both (agrammatic and paragrammatic). All raters were experienced language scientists, with a mix of backgrounds in stroke and aphasia research, speech-language pathology, and linguistics.

paragrammatic errors were classified as those not resulting from an overall reduction or omission of function words/morphemes or structures, but rather grammatical errors with a general presence of functional elements. Using these criteria, individual paragrammatic errors could be omissions. For instance, the utterance “…and they’re **visit**” is ungrammatical because of the omission of the progressive morpheme “-ing”. However, this subject did not appear to omit inflections generally (indeed, the progressive auxiliary “are” is present in the verb contraction in the same utterance), therefore such utterances were taken as evidence for paragrammatism rather than agrammatism. agrammatic subjects by contrast were defined as having an overall deficit of functional word and morpheme omission and reduced sentence complexity.

Of the 100 original subjects, we successfully rated 53, and these were included in the following analyses. Forty-seven subjects were unable to be rated due to audio quality issues (N=2) or severely limited speech output or reduced intelligibility as a result of concomitant motor speech problems (apraxia of speech and/or dysarthria) or severe aphasia (N=45). Of the 53 successfully classified subjects, 21 were assigned to the parparagrammatic group, 11 to the agrammatic group, and 17 to the NGD group. Four subjects were classified as both (agrammatic and paragrammatic) based on the presence of both features in these subjects’ speech. Table 1 provides demographic data for each classified group. While Broca’s aphasia was closely associated with agrammatism, and Wernicke’s and conduction aphasia were closely associated with paragrammatism, a variety of aphasia subtypes were classified within each grammatical category. As lesion volume was significantly larger in the agrammatic group, all of our lesion-symptom mapping analyses included lesion volume as a covariate.

**Table 1.** Demographic information for each group of subjects.

|  | Number | Sex | Age<br>(years) | Months<br>post-stroke | Education<br>(years) | Number<br>w/AOS | WAB<br>AQ | Lesion<br>volume<br>(cm<sup>3</sup>) | Aphasia<br>Diagnosis |
| --- | --- | --- | --- | --- | --- | --- | --- | --- | --- |
| PARAGRAMMATIC | 25 | 9 Female | 62.16*<br>(9.91) | 43.52*<br>(36.42) | 15.04^<br>(2.25) | 8* | 63.50^<br>(17.51) | 108.78*<br>(74.53) | 3 anomia, 6 Broca's, 13 conduction, 3 Wernicke's |
| AGRAMMATIC | 15 | 6 Female | 51.40*^<br>(13.60) | 79.07*<br>(56.09) | 16.07<br>(2.89) | 12*^ | 61.41^<br>(12.92) | 212.66*^<br>(55.11) | 1 anomia, 12 Broca's, 1 conduction, 1 global |
| NGD | 17 | 4 Female | 61.06<br>(12.71) | 66.71<br>(76.51) | 16.41<br>(2.00) | 7 | 80.28<br>(14.02) | 80.66<br>(73.81) | 10 anomia, 6 Broca's, 1 conduction |
| BOTH | 4 | 2 Female | 50.75<br>(15.90) | 91.00<br>(47.94) | 16.50<br>(3.42) | 2 | 65.18<br>(12.51) | 150.64<br>(38.92) | 3 Broca's<br>1<br>conduction |

\*significant difference ( $p < 0.05$ ) between the PARAGRAMMATIC and AGRAMMATIC groups.
^significant difference between PARAGRAMMATIC/AGRAMMATIC and NO GRAMMATICAL DEFICIT.
NGD = NO GRAMMATICAL DEFICIT; WAB AQ = performance on the Western Aphasia Battery summary measure for overall aphasia severity. Four subjects classified as BOTH (i.e., showing features of both AGRAMMATISM and PARAGRAMMATISM) were included in the AGRAMMATIC and PARAGRAMMATIC groups shown here, as well as separately.

### Examples of agrammatic errors (in subjects classified as agrammatic only)

Subject 127: “Cinderella very dress up”

Subject 127: “Cinderella one shoe”

Subject 129: “two girls and boy bad”

Subject 183: “daughters angry and uh sad”

Subject 183: “Cinder[ella]… poor…and no rooms”

Subject 1040: “fairy… um… Cinderella dress pretty… far away kinda… uh…the wish upon a star… um… the horse pretty and… “

Subject 1040: “Cinderella all dressed and… slippers”

Subject 171: “a man… slippers.. and.. look…”

Subject 171: “and slipper… fit”

Subject 171: “and… crown… and… married”

### Examples of parparagrammatic errors (in subjects classified as parparagrammatic only)

Subject 1034: “to **live his father and stepmother**” – verb complement selection violation

Subject 1034: “three other **sister**” – number agreement error

Subject 1034: “he had to do **all the works**” – incorrect plural inflection of mass noun

Subject 1034: “wanted to **make a trick her**” – fusion error: *trick* is both a noun and a verb

Subject 1041: “she **came the house**” - verb complement selection violation

Subject 1007: “she **goed** back to the boy” - use of regular past tense inflection for irregular verb

Subject 1010: “**tooked** her dress” - use of regular past tense inflection in addition to irregular past tense formation

Subject 170: “one stepfather and **a stepchildren**” – number agreement error

Subject 184: “and she got **the flied**” – verb in noun position; regular past tense inflection on irregular verb

Subject 194: “all they want to do is look **the** pretty” – article inserted incorrectly

Subject 194: “one of her slippers that **were** glass” – number agreement error

Subject 194: “I want to find this girl that I saw **you**” – pronoun inserted into gap of relative clause

Subject 194: “she ran **the steps**” – verb complement selection violation

Subject 198: “she **was** met the prince” – incorrect auxiliary

Subject 198: “she turned out **to** the same person” – dropping of copula from infinitive

Subject 198: “**the some** happen to them” – multiple articles without noun

Subject 201: “this is not what **good**” – unclear construction, perhaps omission of auxiliary (“what was good” -> “what good”)

Subject 201: “had something was going **to back**” – strange construction; omission of article inside prepositional phrase (to *the* back -> to back)

Subject 209: “two women **is** ugly” – number agreement error

Subject 209: “two ugly **child was**” – number agreement error(s)

Subject 209: “two **man**” – number agreement error

Subject 209: “the queen and king **is** there” – number agreement error

Subject 209: “**the** Cinderella is found in the one” – incorrect use of definite article for proper name

Subject 143: “she **flied** around” – use of regular past tense inflection for irregular verb

Subject 143: “and they’re **visit**” – omission of progressive inflection

### Examples errors in subjects classified as both (both agrammatic and paragrammatic)

Subject 1002: “the girl run” –omission of verb inflection

Subject 1002: “And … evil woman… and locked the door” – telegraphic

Subject 1002: “The twins and evil mother” – omission of article on second noun of conjunct

Subject 1002: “It is girls” – agreement error

Subject 1002: “The girl is what can we do” – sentence monster

Subject 1002: “And the prince and the Cinderella is happily forever” – agreement error, incorrect use of definite article for proper name

Subject 1024: “Cinderella cleaning so nice” - telegraphic

Subject 1024: “Cinderella is not going because I think that list” - telegraphic

Subject 1024: “so… different dress and…” - telegraphic

Subject 1024: “but mom is list for cleaning, vacuum, whatever” – telegraphic, perhaps sentence monster

Subject 1024: “Cinderella you need to go [be]fore midnight because it’s broke your, spell your broken” – sentence monster

Subject 1024: “Cinderella and prince is going to marry” – number agreement error

Subject 1044: “Cinderella is good” – simplification of sentence structure

Subject 1044: “slippers in the drawer is no good” – omission of definite determiner, agreement error

Subject 1044: “Cinderella is danced with her maid” – incorrect inflection (substitution of -ed foring)

Subject 190: “There’s girl and boy” – simplification of sentence structure, omission of articles

Subject 190: “The other one, bad…bad…” – omission of copula

Subject 190: “The other one, bad bad” – omission of copula

Subject 190: “Everything nice” – omission of copula

Subject 190: “The girl… he’s really nice” – gender agreement error

Subject 190: “It’s moving the talking” – sentence monster

### Reliability

To determine inter-rater agreement, each of the four expert raters individually re-rated videos from n=10 subjects, i.e., >20% of the sample that could be rated. Fleiss’ kappa was used to determine rater agreement between each of the four raters. There was “good” agreement between each rater, k=0.61, (95% CI., 0.44 to 0.77), p<0.0001. Individual kappa for each of grammatical classification ranged from “fair” (parparagrammatic classification, k=0.25), to “good” (both, no grammatical deficit, k=0.67 for both classifications), and “very good” (agrammatic classification k=0.90). Consensus agreement was also computed demonstrating the extent to which the first consensus discussion helped with rater clarity and reliability for subject classification. Cohen’s Kappa was used to confirm reliability of rater criteria following the consensus agreement meetings. Cohen’s kappa was 0.86, p<0.0001, suggesting strong reliability for consensus rating criteria.

### Speech fluency and speech rate

To avoid a similar conflation of grammatical abilities with ability from overall speech fluency, we incorporated a speech rate dimension, words per minute (WPM), as a covariate in secondary analyses. WPM was calculated based on CHAT/CLAN transcriptions of each subject’s speech during the same Cinderella task that was used for assessment of grammatical deficits (MacWhinney *et al*., 2000). Speech was transcribed by a speech pathology masters student, using audiovisual speech recordings. Speech rate, as measured in words per minute (WPM), is shown in Fig. 1 (left). For comparison, scores from the WAB-R fluency rating scale (Kertesz, 2007) are shown in Fig. 1 (right). For both measures, the agrammatic group (including the subjects categorized as ‘both’) were the least fluent. With respect to the parparagrammatic and no grammatical deficit groups, the two measures diverged. For WPM, the parparagrammatic group was most fluent, while for WAB-R fluency, the no grammatical deficit group was most fluent. This is because the WAB-R fluency scale incorporates both quantity and quality (e.g. grammatical deficits) of speech output, whereas WPM measures speech rate regardless of quality.

**Figure 1.**
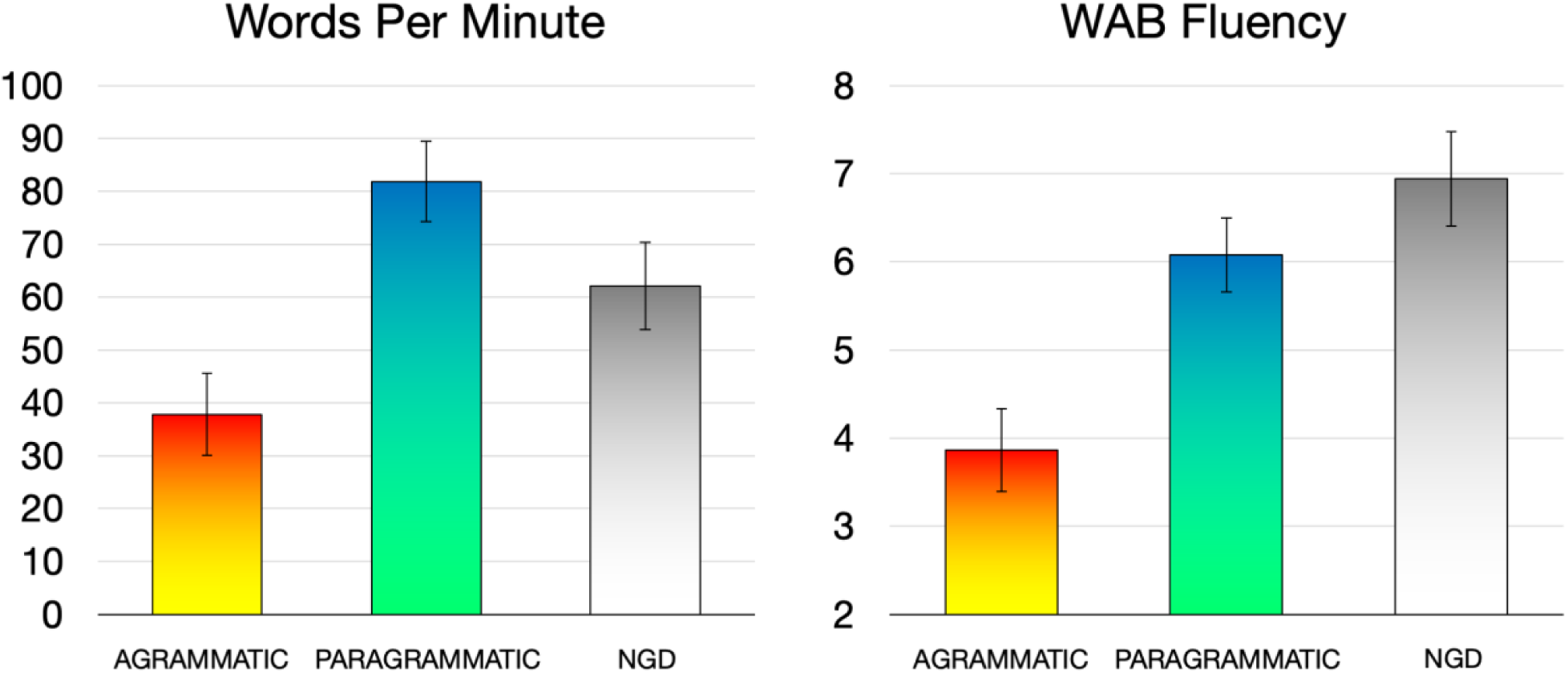
Speech fluency, as measured in words per minute during the Cinderella story (left), and the WAB-R fluency measure (right). Error bars reflect standard error of the mean.

### Neuroimaging Data acquisition and processing

We acquired anatomical MRIs and performed lesion mapping using the same parameters and procedures as described in Fridriksson et al. (2018). Neuroimaging data were collected at the University of South Carolina (MUSC; 15 subjects) and the Medical University of South Carolina (MUSC; 38 subjects). Lesions were demarcated onto each subject’s T2 image by an expert technician (RNN, data from USC) or an expert neurologist (LB, data from MUSC) blind to the behavioral data. Lesion overlap maps for each group included in final analyses are shown in Fig. 2.

**Figure 2.**
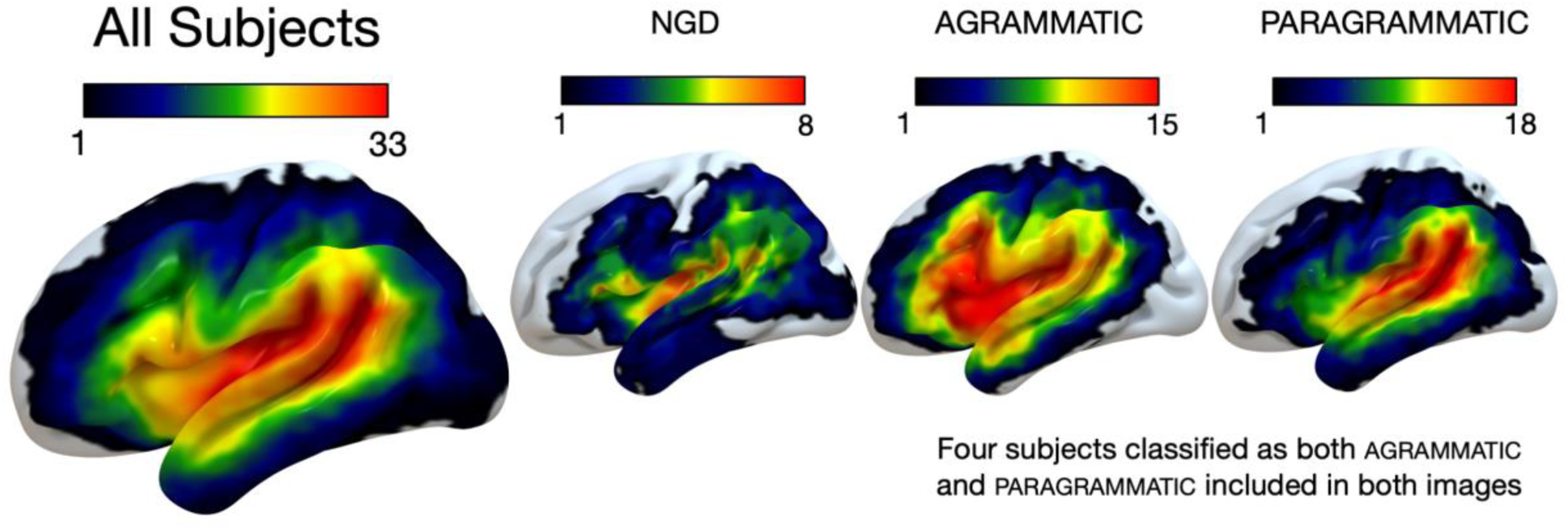
Lesion overlap maps for each group. All Subjects (N=53, max overlap=33), no grammatical deficit (ngd) (N=17, max overlap=8), agrammatic (N=15, max overlap=15), parparagrammatic (N=25, max overlap=18).

### ROI analyses

We created two regions of interest (ROIs) by combining pre-existing parcellations from the Johns Hopkins University atlas (Faria *et al*., 2012). The Broca’s area ROI was created by combining the left inferior frontal gyrus, pars triangularis and pars opercularis. The left posterior superior and middle temporal gyri ROI (pSTG/MTG) was created by combining the posterior superior temporal gyrus and middle temporal gyrus. For each subject, we calculated percent damage to each ROI, and then calculated residual percent damage values after performing a linear regression in SPSS with (i) lesion volume only as a covariate, and (ii) lesion volume and speech rate (WPM) as covariates. We then entered these residual values into independent samples *t*-tests in SPSS, evaluating the effect of agrammatism (subjects classified as agrammatic vs. subjects not classified as agrammatic) and paragrammatism (subjects classified as parparagrammatic vs. subjects not classified as paragrammatic) within each ROI.

In order to ensure that lesion volume was not distributed differently between the agrammatic and parparagrammatic groups within these ROIs, we calculated Levine’s statistic testing for homogeneity of variances between these groups (based on median percent damage, Brown-Forsythe test). The homogeneity of variance was not shown to be different between the agrammatic and parparagrammatic groups within either the Broca’s area ROI (*W*(1,38) = 1.567, p = 0.218) or the pSTG/MTG ROI (*W* (1,38) = 0.027, p = 0.869). Thus, our use of the lesion volume covariate did not appear to interact differently with the distinction between agrammatism and paragrammatism within these ROIs. We also show ROI data without lesion volume covariates in the Supplementary Materials (Supplementary Figure 1).

### Whole-brain analyses

We performed corresponding whole-brain analyses in NiiStat (https://www.nitrc.org/projects/niistat/), using binary logistic regression to remove effects of (i) lesion volume only and (ii) lesion volume and speech rate (WPM) from the behavioral classifications. We then performed one-tailed *t*-tests, using an uncorrected individual voxel-wise threshold of p < 0.05. Only voxels that were damaged in at least 5 subjects (∼10% of sample) were included in the analyses. We also show uncorrected whole-brain results using a stricter statistical threshold, p < 0.001, in the Supplementary Materials (Supplementary Figure 3).

In order to ensure that lesion volume across the whole brain was not distributed differently between the agrammatic and parparagrammatic groups, we calculated Levine’s statistic testing for homogeneity of variances between these groups (based on median percent damage, Brown-Forsythe test). The homogeneity of variance of whole brain lesion volume was not shown to be different between the agrammatic and parparagrammatic groups (*W*(1,38) = 1.851, p = 0.182). Thus, our use of the lesion volume covariate did not appear to interact differently with the distinction between agrammatism and paragrammatism at the whole brain level. We also show whole-brain results without lesion volume covariates in the Supplementary Materials (Supplementary Figure 2).

## Results

Fig. 3 shows results from the ROI analyses. The results revealed a clear double dissociation: when including only lesion volume as a covariate, agrammatism was significantly associated with damage to Broca’s area (t(51) = 3.133, p = 0.001), but not pSTG/MTG (t(51) = -1.152, p = 0.873), while paragrammatism was significantly associated with damage to pSTG/MTG (t(51) = 3.674, p < 0.001) but not Broca’s area (t(51) = -1.499, p.= 0.930). These results also held when adding speech rate as a covariate: agrammatism was significantly associated with damage to Broca’s area (t(51) = 2.458, p = 0.009) but not pSTG/MTG (t(51) = -0.528, p = 0.700), while paragrammatism was significantly associated with damage to pSTG/MTG (t(51) = 2.786, p = 0.004) but not Broca’s area (t(51) = -0.744, p = 0.770).

**Figure 3.**
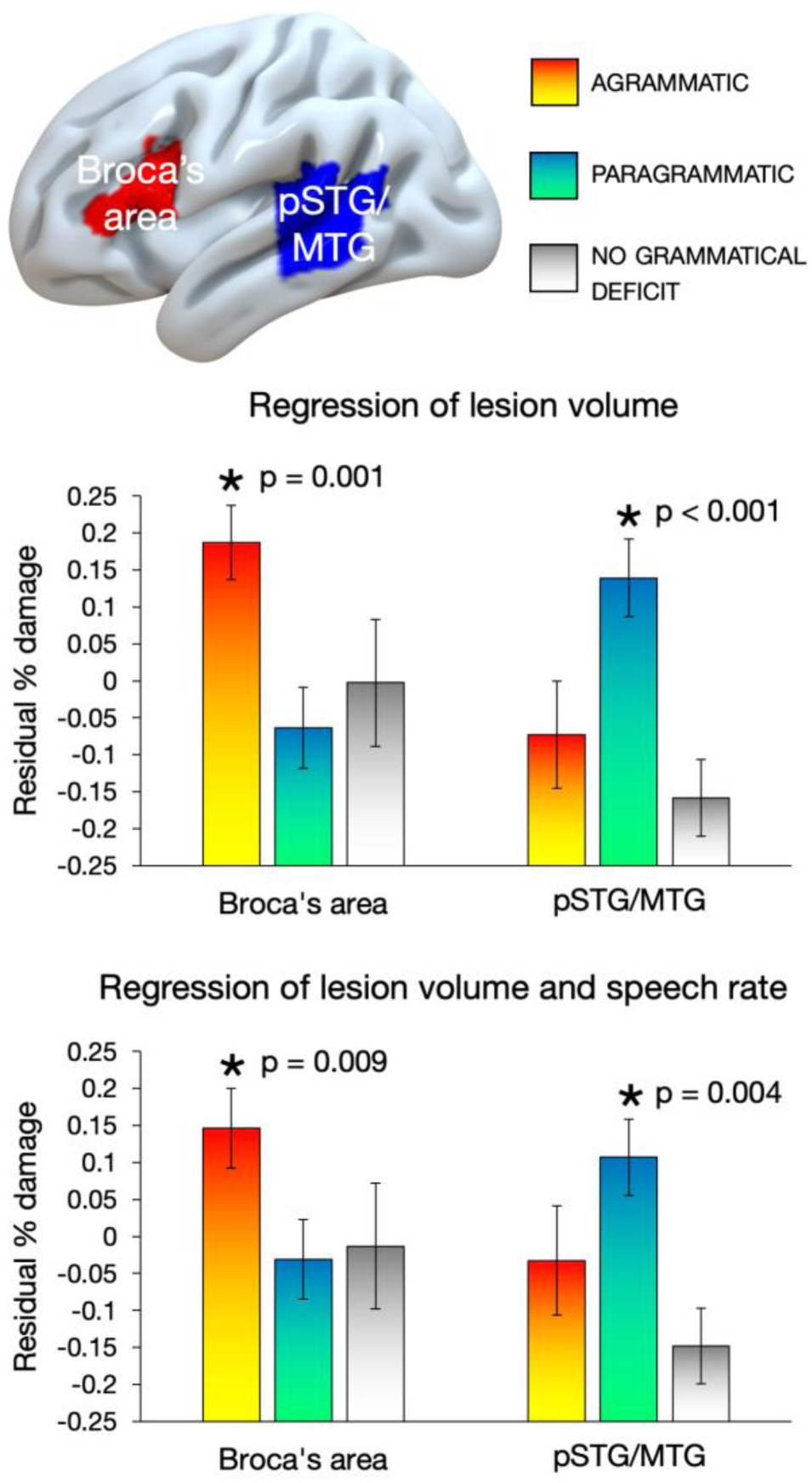
Double dissociation between the effect of agrammatism and the effect of paragrammatism with respect to damage in Broca’s area and the posterior superior and middle temporal gyri (pSTG/MTG) regions of interest (ROIs). TOP: selected ROIs. Residual values of average percent damage for each group to each ROI, after regressing out lesion volume (LV) (MIDDLE) and after regressing out both LV and words per minute (WPM) (BOTTOM) using linear regression in SPSS. Error bars reflect standard error of the mean. Asterisks indicate significant effects when performing a Bonferroni correction for multiple comparisons and an alpha of 0.0125. pSTG/MTG: posterior superior and middle temporal gyri. Four subjects classified as both (both agrammatic and paragrammatic) were included in both analyses. In the Supplementary Materials, we show corresponding data without the lesion volume covariate (Supplementary Figure 1).

Fig. 4 shows the results of our whole-brain analyses (local maxima peak coordinates are shown in Table 2). With respect to the effect of agrammatism (Fig. 4, hot colors), both analyses (with and without regression of WPM) revealed damage to the IFG and middle frontal gyrus, encroaching into the insula, with minor damage to the anterior supramarginal gyrus (SMG). When only including lesion volume as a covariate, agrammatism was also associated with damage to left ventral precentral gyrus, but these effects were not present when speech rate was included as a covariate. paragrammatism (Fig. 4, cool colors) was associated with damage to left posterior temporal and inferior parietal cortex, centered on the posterior superior temporal gyrus and middle temporal gyrus, including the inferior and posterior parts of the SMG. Overall, regressing out the effect of speech rate (WPM) reduced the strength and the spatial extent of the damage associated with agrammatism and paragrammatism while maintaining essentially the same spatial location. There was no spatial overlap in the damage associated with each type of grammatical deficit, whether or not WPM was included as a covariate.

**Table 2.** Local peak coordinates for the uncorrected whole-brain analyses (all coordinates in left hemisphere).

| Region | X | Y | Z | Z-value |
| --- | --- | --- | --- | --- |
| *PARAGRAMMATISM (regressing out the effect of lesion volume only)* |  |  |  |  |
| STG/STS (posterior) | -47 | -45 | 11 | 3.97 |
| Temporal-parietal junction | -45 | -50 | 19 | 4.05 |
| Inferior parietal lobule/angular gyrus | -46 | -53 | 32 | 4.38 |
| MTG/STS (middle-posterior) | -43 | -39 | 2 | 4.15 |
| STG (middle) | -51 | -19 | 5 | 3.86 |
| *PARAGRAMMATISM (regressing out the effect of lesion volume and WPM)* |  |  |  |  |
| Temporal-parietal junction | -42 | -51 | 20 | 3.59 |
| Superior temporal gyrus | -47 | -27 | 8 | 3.00 |
| Supramarginal gyrus | -55 | -48 | 31 | 3.20 |
| Middle temporal gyrus | -50 | -41 | -1 | 3.14 |
| Superior temporal gyrus | -57 | -37 | 12 | 3.41 |
| *AGRAMMATISM (regressing out the effect of lesion volume only)* |  |  |  |  |
| Postcentral gyrus | -67 | -10 | 22 | 3.98 |
| Inferior frontal gyrus (pars triangularis) | -53 | 22 | 26 | 5.22 |
| Inferior frontal gyrus (pars triangularis)/middle frontal gyrus | -36 | 23 | 31 | 4.85 |
| *AGRAMMATISM (regressing out the effect of lesion volume and WPM)* |  |  |  |  |
| Inferior frontal gyrus (pars triangularis) | -53 | 22 | 26 | 4.74 |
| Inferior frontal gyrus (pars triangularis)/middle frontal gyrus | -45 | 28 | 29 | 4.50 |

**Figure 4.**
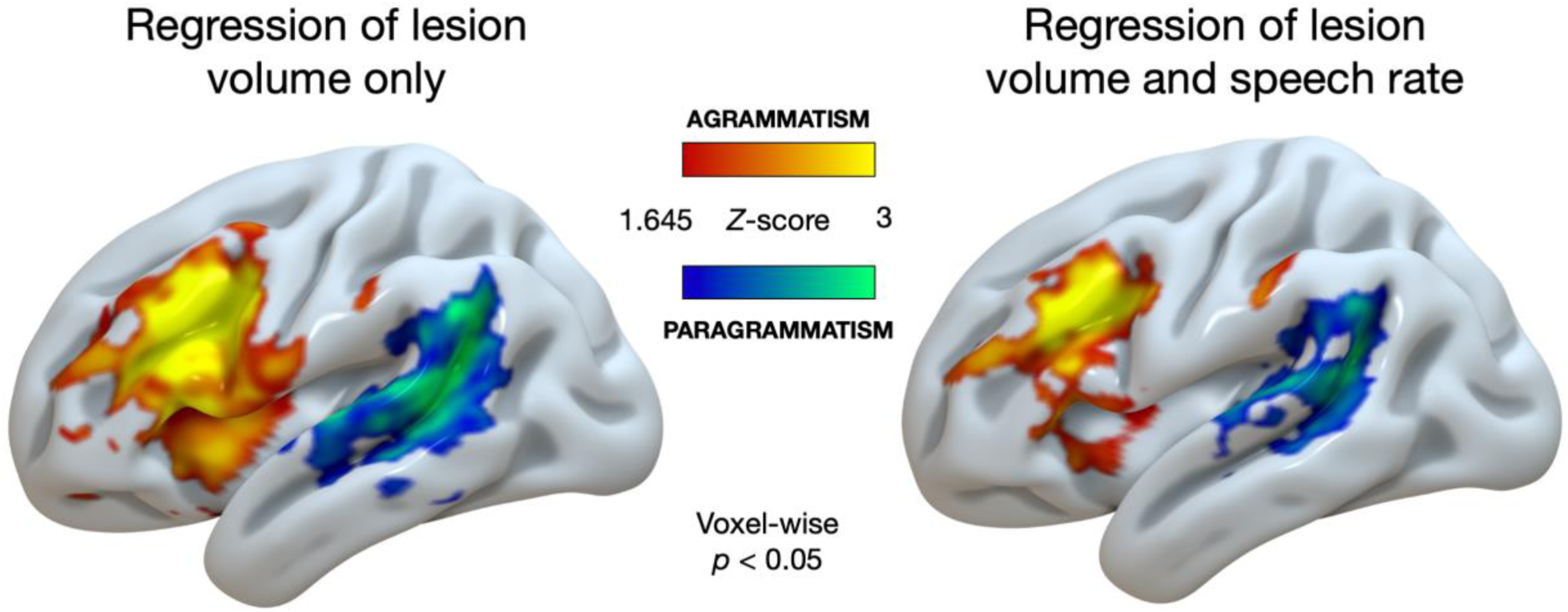
Uncorrected whole-brain analyses (voxel-wise p < 0.05) of the effects of agrammatism (red-yellow) and paragrammatism (blue-green) displayed on the cortical surface an inflated left hemisphere brain template in MNI space. Left: only lesion volume included as a covariate. Right: both lesion volume and speech rate (words per minute) included as covariates. agrammatism = effect of agrammatism (agrammatic, including both > no grammatical deficit and paragrammatic, excluding both), paragrammatism = effect of paragrammatism (paragrammatic, including both > no grammatical deficit and agrammatic, excluding both).

In the Supplementary Materials, we show corresponding results without the lesion volume covariate (Supplementary Figure 2). We also show uncorrected whole-brain analyses at a stricter statistical threshold (p < 0.001) (Supplementary Figure 3).

To identify the relation between damage associated with agrammatism and impaired speech rate, we performed uncorrected whole-brain analyses comparing the effects of reduced WPM and agrammatism (Fig. 5). Reduced WPM was associated with damage to left inferior post-central gyrus, precentral gyrus, middle frontal gyrus, and inferior frontal gyrus. There was notable overlap between the lesion distribution associated with reduced WPM and agrammatism in the left precentral gyrus and inferior frontal gyrus when only regressing lesion volume. However, regressing out the effect of WPM from agrammatism, and removing agrammatic subjects from the analysis of reduced WPM (i.e., only analyzing the effect of reduced WPM in patients without agrammatism, n = 38), revealed minimal overlap between agrammatism and reduced WPM. Namely, reduced WPM in absence of agrammatism was associated with damage to left inferior precentral gyrus, and agrammatism (covarying out WPM) was associated with damage to left inferior and middle frontal gyrus.

**Figure 5.**
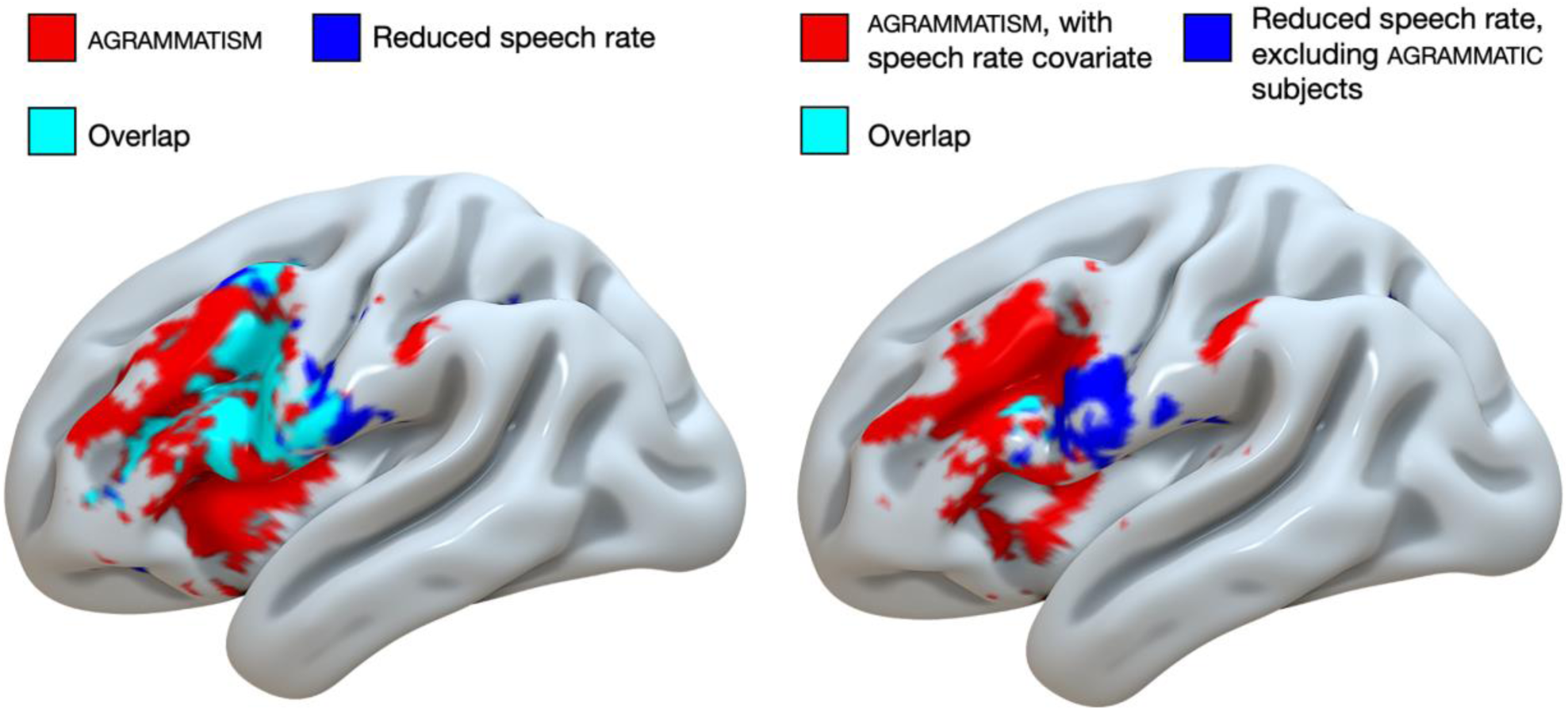
Uncorrected whole-brain analyses (voxel-wise p < 0.05) of the effects of agrammatism (red), reduced speech rate in words per minute (blue), and their overlap (cyan) displayed on the cortical surface an inflated left hemisphere brain template in MNI space. Left: only lesion volume included as a covariate. Right: speech rate included as an additional covariate in the analysis of agrammatism (red), and excluding agrammatic subjects in the analysis of reduced speech rate (blue, n = 38), with minimal overlap (cyan).

## Discussion

Our results revealed a clear double dissociation: agrammatic speech (but not parparagrammatic speech) is significantly associated with damage to Broca’s area, while parparagrammatic speech (but not agrammatic speech) is significantly associated with damage to the left posterior superior and middle temporal gyri (pSTG/MTG). This double dissociation held even when accounting for variability in speech rate (words per minute). The association we identified between agrammatism and inferior and middle frontal damage is consistent with previous lesion-deficit mapping studies (Sapolsky *et al*., 2010; Wilson *et al*., 2010; den Ouden *et al*., 2019), and the association we observed between damage to left posterior temporal-parietal cortex with paragrammatism is consistent with previous case studies (Yagata *et al*., 2017; Wilson *et al*., 2018*a*; *b*). The association of paragrammatism with posterior damage is also consistent with the general association of paragrammatism with fluent aphasia, which typically involves damage to left posterior temporal-parietal systems (Buchsbaum *et al*., 2011; Ogar *et al*., 2011), although we note that patients from both fluent and non-fluent aphasias were present in both our agrammatic and parparagrammatic groups.

There has been strikingly little focus on understanding the nature of paragrammatism and its neural correlates, in contrast to agrammatic speech, which has had a much greater focus (Kean, 1985). This is in part due to the strong historical association of Broca’s aphasia and agrammatism as reflecting a core syntactic deficit (Caramazza and Zurif, 1976; Schwartz *et al*., 1980; Grodzinsky, 1986; 2000), and a corresponding focus on the role of the left IFG in syntax (Friederici, 2017; Grodzinsky and Santi, 2008). Our results drive home the importance of considering paragrammatism as an independent grammatical disorder, resulting from damage to distinct brain systems from those implicated in agrammatic speech.

The limited attention that has been paid to paragrammatism in past research has often been in the service of the hypothesis that agrammatism and paragrammatism are different degrees of adaptation to a common underlying deficit (Heeschen, 1985; Heeschen and Kolk, 1988; Bates and Wulfeck, 1989; Kolk and Heeschen, 1992; Bates and Goodman, 1997), including a critique of the supposed unique relation between Broca’s area damage and agrammatism (Dick *et al*., 2001). This hypothesis is supported by the fact that paragrammatic errors can be observed in non-fluent agrammatic speakers when they forced to produce speech more quickly than their natural pace (Heeschen and Kolk, 1988). However, the double dissociation we identified here, which held even when incorporating words per minute as a covariate, speaks against this hypothesis. Consistent with a more fundamental distinction between agrammatism and paragrammatism, Casilio *et al*. (2018) recently reported a factor analysis for a large range of perceptual measures similarly extracted from connected speech samples (N=24 people with aphasia). They show that paragrammatism is sharply distinguished from agrammatic features such as omission of function words/morphemes. Paragrammatism was associated with a variety of speech deficits such as abandoned utterances, empty speech, semantic paraphasias, phonemic paraphasias, and neologisms. Nevertheless, we do note that many of the paragrammatic errors we identified here (particularly inflectional errors such as agreement mismatch) do not follow straightforwardly from lexical-semantic substitutions and appear to constitute a separate deficit.

An additional reason that previous research has focused largely on agrammatism to the exclusion of paragrammatism is that it is difficult to clearly identify paragrammatism as distinct from these other disturbances to speech output. This is reflected in the relatively modest inter-rater agreement of paragrammatism that our expert raters obtained in this study (k=0.25), whereas the inter-rater agreement for agrammatism was much greater (k=0.90). One potential solution to this issue is to use an objective classification of paragrammatic deficits rather than relying on a perceptual rating of paragrammatism (as in the present study and Casilio et al., 2018). However, we were careful to note that isolated speech samples can be difficult to characterize as agrammatic or paragrammatic using an objective criterion. This is because if paragrammatism reflects a random misuse of grammar, omissions of functional elements will occasionally result, incorrectly appearing to indicate agrammatism. Therefore an important contribution will be to develop clear diagnostic criteria for paragrammatism and to investigate the degree to which paragrammatic deficits can be separated from non-grammatical deficits (such as lexico-semantic errors), including both subjective perceptual and objective criteria.

The pSTG/MTG and Broca’s area have previously been implicated in syntactic processing, with remarkably similar effects in both areas. They both exhibit increased activity for grammatically structured linguistic materials relative to unstructured word lists regardless of the richness of conceptual-semantic content (Pallier *et al*., 2011; Goucha *et al*., 2015; Matchin *et al*., 2017), and damage to both regions has been implicated in deficits comprehending syntactically complex materials (Tyler *et al*., 2011). While these previous studies have shown largely similar syntactic effects in both regions, the double dissociation revealed in the present study supports a functional distinction between them.

Previous lesion-deficit mapping studies of basic sentence and syntactic *comprehension* in the absence of working memory confounds have also primarily identified similar left posterior temporal-parietal areas (Wilson and Saygin, 2004; Thothathiri *et al*., 2012; Pillay *et al*., 2017; Fridriksson *et al*., 2018; Rogalsky *et al*., 2018; den Ouden *et al*., 2019). This suggests a role for left posterior temporal-parietal cortex in grammatical processes that underlie both comprehension and production. Matchin and Hickok (2019) put forward the theory that left posterior middle temporal gyrus underlies hierarchical lexical-syntactic structure, and that sentence comprehension and paragrammatic production deficits follow from difficulties with hierarchical syntax. For example, consider paragrammatic errors such as incorrect subject-verb number agreement, e.g., “the queen and king **is** there”. Subject-verb agreement in English must be calculated over structure. In the paragrammatic example, both “queen” and “king” occur in the singular. However, the correct agreement feature is plural: the verb agrees with the conjoined noun phrase, rather than the individual nouns. Thus, the paragrammatic error can be seen as the lack of constraining hierarchical structural relations.

Our results reinforce previous findings regarding the association of agrammatic production and damage to the left inferior frontal cortex (Sapolsky *et al*., 2010; Wilson *et al*., 2010; Den Ouden *et al*., 2019). Broca’s area has long been associated with speech production, and most current research supports a role for higher-level planning of language rather than lower-level motor execution (Basilakos *et al*., 2015; Flinker *et al*., 2015). Matchin and Hickok (2019) suggest that the anterior portion of Broca’s area, the pars triangularis, supports production via morpho-syntactic sequences that are necessary for converting hierarchical structures into linear speech output, and that when damaged, speech becomes agrammatic.

On this point, we note that while previous research has found a general overlap between effects of reduced speech rate and agrammatism (Wilson *et al*., 2010), the effect of agrammatism we observed in left inferior and middle frontal cortex remained even when regressing out the effect of speech rate (albeit somewhat weakened). Additionally, reduced speech rate in subjects without agrammatism was associated with a distinct non-overlapping cluster in inferior precentral gyrus. Finally, a previous lesion-symptom mapping study distinguishing between aphasia and apraxia of speech found that these syndromes strongly dissociate (Basilakos et al., 2015). In that study, focusing on frontal cortex, apraxia of speech was associated with damage to inferior precentral gyrus, whereas aphasia was associated with damage to anterior inferior frontal gyrus (pars triangularis and pars orbitalis). In the present study, a large proportion of the agrammatic speakers were indeed also diagnosed with apraxia of speech (12/15, versus 8/25 of the parparagrammatic speakers). These previous results, however, including our regression analysis, show that apraxia of speech cannot be the sole driving force behind agrammatism and it is more likely that the two co-occur due to the anatomical proximity of their neural substrates, even if these do not overlap.

Neither agrammatism nor paragrammatism was associated with damage to the left anterior temporal lobe (ATL) in our whole-brain analyses. The finding agrees with previous lesion-symptom mapping studies of agrammatic production failing to identify effects in the ATL (Sapolsky *et al*., 2010; Wilson *et al*., 2010; den Ouden *et al*., 2019). However, some previous studies have reported associations between non-canonical sentence comprehension deficits and damage to the left ATL in both chronic and acute stroke that might imply a syntactic function of this region (Dronkers *et al*., 2004; Magnusdottir *et al*., 2013). We note that both of these studies did not control for lesion volume. As reported in Supplementary materials (Supplementary Figure 2), we show that agrammatism was in fact significantly associated with damage to the ATL when lesion volume was not included as a covariate. Given this, we suggest that stroke-based lesion-symptom mapping studies of noncanonical sentence comprehension that control for lesion volume will identify effects primarily in left posterior temporal areas and not the ATL, as reported by Rogalsky et al (2018).

Although we started with a large sample (100 subjects), the difficulty in obtaining enough speech output in the experimental task combined with the partitioning of subjects into distinct group resulted in less overall statistical power. Omitting subjects with low speech output also likely precludes analysis of patients with both severe agrammatic and nonfluent speech. Thus, patients who might be highly informative to our lesion-symptom mapping results are precluded by the nature of our approach: analyzing production during spontaneous discourse/narration. These issues should be addressed in future studies.

Overall, the double dissociation identified here confirms the predictions of Matchin and Hickok (2019): agrammatism (but not paragrammatism) follows from inferior frontal damage, while paragrammatism (but not agrammatism) follows from posterior temporal damage. This suggests that agrammatism and paragrammatism are distinct syndromes as opposed to reflecting alternative adaptations to the same underlying grammatical deficit (Kolk and Heeschen, 1992; Dick *et al*., 2001). Both basic research into the neural organization of language as well as research into the nature of aphasia symptoms and recovery will benefit from increased focus on paragrammatism itself and its distinction from agrammatism.

### Key Term Definitions

#### Agrammatism

Common symptom of non-fluent aphasia. Syntax is simplified and function words and morphemes (e.g. articles, past tense) are omitted.

#### Paragrammatism

Common symptom of fluent aphasia. Syntactic errors not attributable to overall simplification or omission of functional elements (e.g. agreement errors).

#### Lesion-Symptom Mapping

Method of determining the relationship between the extent and location of brain damage with behavioral measures or deficits.

#### Fluency

Ease of production, particularly applied to aphasia. Includes both quantity and quality of speech.

#### Fluent Aphasia

Aphasias characterized by typically fluent production, including Wernicke’s aphasia, conduction aphasia, and anomia.

#### Non-fluent Aphasia

Aphasias characterized by typically non-fluent production, including Broca’s aphasia, global aphasia, and anomia.

#### Western Aphasia Battery (WAB)

Battery of tests to assess language abilities. Performance is used to identify and classify aphasia in people with neurological disfunction.

#### Syntax

Sentence structure. Includes both linear information (such as word order) and hierarchical information (grouping of words into phrases).

#### Morpho-syntax

The internal structure of words and sentences. Includes syntactic features marked on words (e.g., plural *-s*, past tense *-ed*,).

#### Agreement

Dependency between two linguistic elements. E.g. subject-verb agreement in English, where *-s* on a verb indicates a third person/singular subject.

## Acknowledgements

We would like to thank Grigori Yourganov, Roger Newman-Norlund, Chris Rorden, and Leo Bonilha for their technical assistance. We would also like to thank Leigh Ann Spell, Allison Croxton, Anna Doyle, Michele Martin, Katie Murphy and Sara Sayers for their assistance with data collection. We would like to thank graduate student clinicians in the Aphasia Lab for transcribing and coding speech samples.

