## Supplementary Materials for "Agrammatism and paragrammatism: a cortical double dissociation revealed by lesion-symptom mapping"

The large discrepancy between the AGRAMMATIC and PARAGRAMMATIC groups with respect to lesion volume (average lesion volume in AGRAMMATIC compared to PARAGRAMMATIC), with no significant differences in the distribution of lesion volume (see main text), justified our inclusion of lesion volume as a secondary covariate of no interest in all of our analyses. In Supplementary Figures 1 and 2, we show our data without lesion volume as a covariate.

Our ROI analyses revealed that AGRAMMATISM was very strongly associated with damage to Broca's area, even more strongly than without the lesion volume covariate. However, without the lesion volume covariate, AGRAMMATISM was also associated with damage to the posterior superior temporal gyrus/middle temporal gyrus pSTG/MTG. As in the analyses that included the lesion volume covariate, PARAGRAMMATISM was associated with damage to pSTG/MTG but not Broca's area, similar with or without the speech rate covariate.

With respect to our whole brain analyses, AGRAMMATISM was strongly associated with damage to the frontal lobe, encroaching onto the superior temporal lobe: primarily superior temporal gyrus, including the anterior temporal lobe (Supplementary Figure 2, top left). Additional effects in the angular gyrus/posterior MTG were revealed when speech rate was included as a covariate (Supplementary Figure 2, top right). This supports our assertion (from the main text) that anterior temporal damage associated with grammatical deficits may be due to lesion size confounds. The whole-brain effects of PARAGRAMMATISM were in general highly similar with or without the lesion volume covariate, albeit weaker without, centered on posterior STS and encroaching onto posterior STG, MTG, and supramarginal gyrus (SMG) (Supplementary Figure 2, bottom).

Supplementary Figure 3 illustrates our analyses as reported in the main manuscript at a stricter statistical threshold ($p < 0.001$). With respect to our analyses of AGRAMMATISM, significant voxels were revealed in the inferior frontal gyrus (IFG; pars triangularis and pars opercularis) with a larger cluster in posterior inferior frontal sulcus/middle frontal gyrus (pIFS/MFG). Similar results obtained when the speech rate covariate was added, although the significant cluster was more restricted to the pIFS/MFG, with minimal overlap on superior IFG (pars triangularis and pars opercularis). With respect to our analyses of PARAGRAMMATISM, when only the lesion volume covariate was added, significant voxels were revealed in posterior STG,

STS, and supramarginal gyrus (SMG). When the speech rate covariate was added, one cluster in posterior STG/SMG was significant (274 voxels).

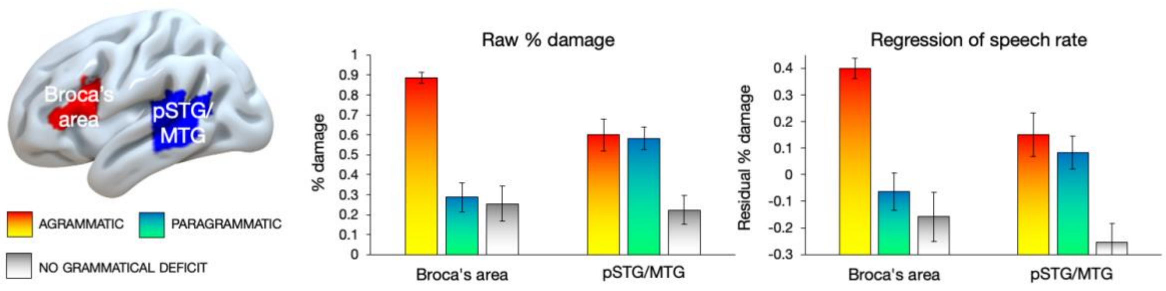


Supplementary Figure 1. ROI percent damage plots without lesion volume covariate.
LEFT: selected ROIs. Average percent damage for each group to each ROI, with no covariates (MIDDLE) and after regressing out words per minute (WPM) (RIGHT) using linear regression in SPSS. Error bars reflect standard error of the mean. pSTS/MTG: posterior superior and middle temporal gyri. Four subjects classified as BOTH (AGRAMMATIC and PARAGRAMMATIC) were included in both analyses.

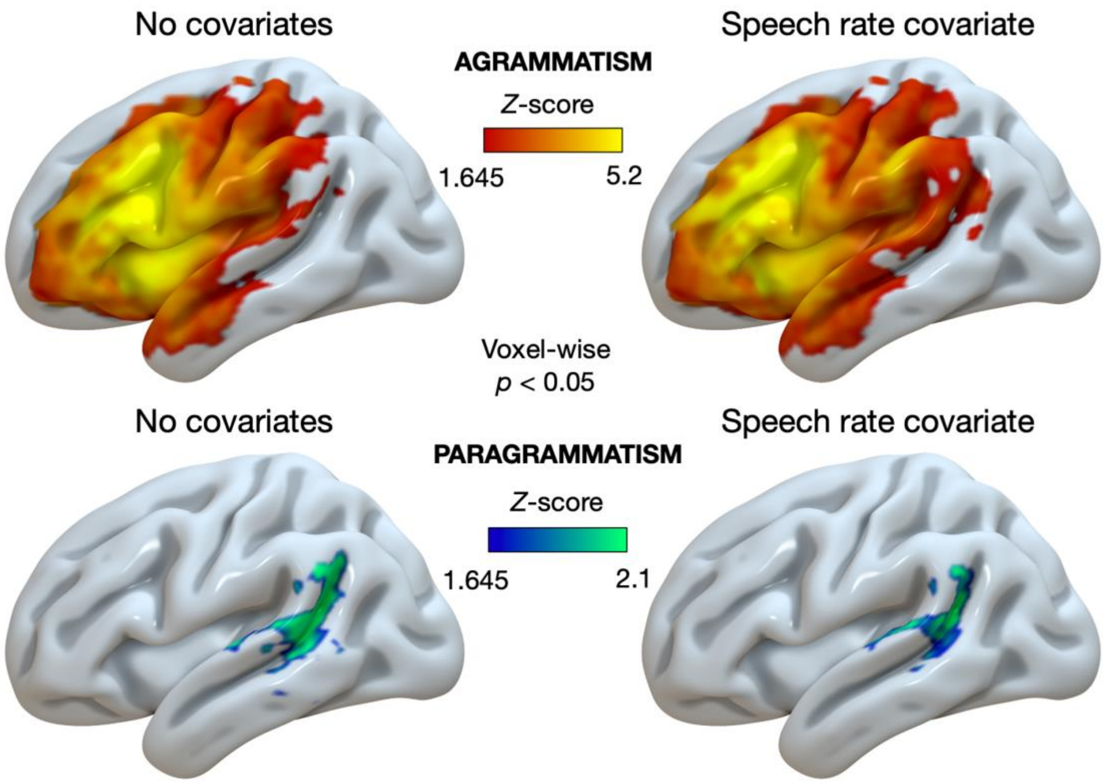

Supplementary Figure 2. Uncorrected whole-brain analyses without the lesion volume covariate (voxel-wise $p < 0.05$) of the effects of AGRAMMATISM (red-yellow) and PARAGRAMMATISM (blue-green) displayed on the cortical surface an inflated left hemisphere brain template in MNI space. LEFT: no covariates. RIGHT: only speech rate (words per minute) included as a covariate. AGRAMMATISM = effect of AGRAMMATISM (AGRAMMATIC, including BOTH > NO GRAMMATICAL DEFICIT and PARAGRAMMATIC, excluding BOTH), PARAGRAMMATISM = effect of PARAGRAMMATISM (PARAGRAMMATIC, including BOTH > NO GRAMMATICAL DEFICIT and AGRAMMATIC, excluding BOTH). Note that different scales are used for the analyses of AGRAMMATISM and PARAGRAMMATISM, given the differences in the strength of the effects without the lesion volume covariate.

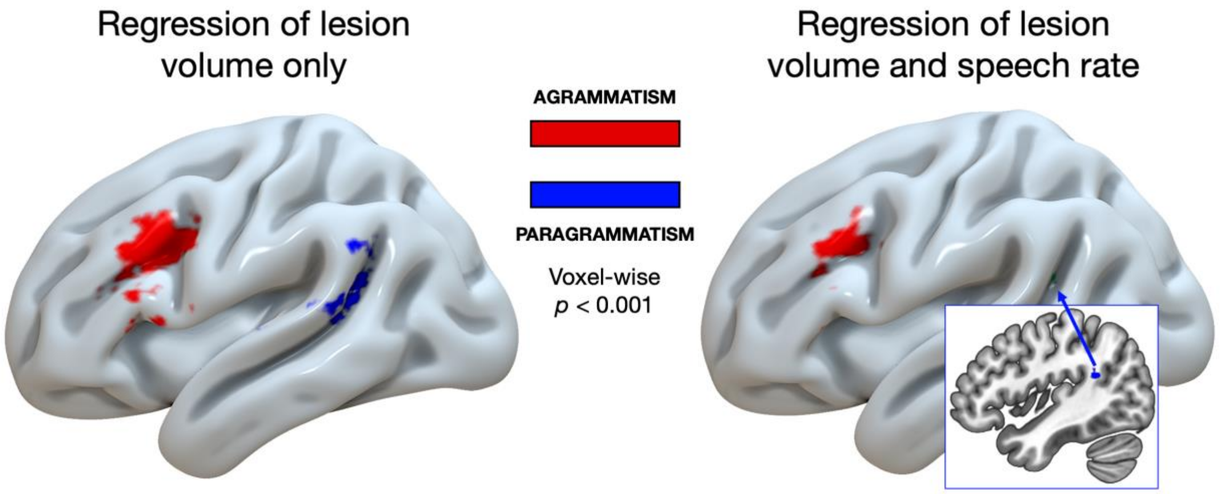


Supplementary Figure 3. Uncorrected whole-brain analyses at a stricter statistical threshold (voxel-wise $p < 0.001$) of the effects of AGRAMMATISM (red) and PARAGRAMMATISM (blue) displayed on the cortical surface an inflated left hemisphere brain template in MNI space. LEFT: only lesion volume included as a covariate. RIGHT: both lesion volume and speech rate (words per minute) included as covariates. AGRAMMATISM = effect of AGRAMMATISM (AGRAMMATIC, including BOTH > NO GRAMMATICAL DEFICIT and PARAGRAMMATIC, excluding BOTH), PARAGRAMMATISM = effect of PARAGRAMMATISM (PARAGRAMMATIC, including BOTH > NO GRAMMATICAL DEFICIT and AGRAMMATIC, excluding BOTH). Inset indicates the location of the significant cluster of voxels for the analysis of PARAGRAMMATISM including both lesion volume and speech rate as covariates. Note that this cluster included 274 significant voxels partially overlapping in white matter, which were obscured when warping to the cortical surface for display purposes.
